# Hippocampal-like network dynamics underlie avian sharp wave-ripples

**DOI:** 10.1101/825075

**Authors:** Hamed Yeganegi, Harald Luksch, Janie M. Ondracek

## Abstract

Sharp wave ripples (SWR) represent one of the most synchronous population patterns in the mammalian brain. Although SWRs are highly conserved throughout mammalian evolution, the existence of SWRs in non-mammalian species remains controversial. We reexamined the existence of avian SWRs by recording the brain activity during sleep and under anesthesia in two species of birds, the zebra finch and the chicken. Electrophysiological recordings using silicon probes implanted in the avian telencephalon revealed highly dynamic switching between high and low delta phases during sleep. High delta phases were composed of large-amplitude, negative deflections (sharp waves) that coincided with a high frequency oscillation (ripple). Correlation analysis revealed that these events were highly synchronous and spanned a large anatomical range of the avian telencephalon. Finally, detailed spike analysis revealed that an increase in the population spiking activity coincided with the occurrence of SWRs, that this spiking activity occurred in specific sequences of spike patterns locked to the SWRs, and that the mean population spiking activity peaked prior to the trough of the negative deflection. These results provide the first evidence of avian SWRs during natural sleep and under anesthesia, and suggest that the evolutionary origin of SWR activity may precede the mammalian-sauropsid bifurcation.

## Introduction

Hippocampal sharp wave-ripples (SWR) represent the most synchronous population pattern in the mammalian brain. Cornelius Vanderwold first observed SWRs in rats in 1969 (1), and John O’Keefe later investigated this hippocampal activity in reference to the spatial memory of rats (2). Since then, SWRs have been observed in the hippocampus of every mammal investigated so far, including humans (3).

SWRs are a complex of two distinct field potentials that occur in the hippocampus: a sharp wave and a ripple. Sharp waves occur during slow wave sleep (SWS) and quiet wakefulness, and these large amplitude, negative deflections reflect massive excitation of neurons in CA1 by CA3 pyramidal neurons (4). Sharp waves are often - although not always - associated with a fast oscillatory pattern of the local field potential (LFP) in the pyramidal layer of CA1, known as “ripples” (5, 6).

SWRs have a number of important features. First, they are *emergent population events* (3) resulting from the coordinated firing of many neurons. Second, SWRs are associated with enhanced transient excitability in the hippocampus, which gives rise to the large synchrony observed during SWRs. Finally, the spiking activity associated with SWRs is highly structured across neurons, which in mammals reflects a temporally compressed version of sequential neuronal firing patterns experienced by the awake animal (11). These features led to the hypothesis that new memories are formed in a “two-step” process, where novel information is encoded in the awake state and then subsequently consolidated during sleep (12).

Although SWRs appear to be highly conserved through-out mammalian evolution, the existence of SWRs in non-mammalian species remains controversial. Recent work in non-avian reptiles such as the Australian bearded dragon (13, 14) revealed that reptilian SWRs are present during bouts of SWS. Importantly, reptilian SWRs were not recorded in the reptilian hippocampal homologue (the medial cortex), but rather in a brain area known as the dorsal ventricular ridge (DVR). The DVR of reptiles is a large pallial region dorsal to the basal ganglia and homologous to the mammalian amygdala (15).

Evidence for SWRs in the avian brain is much more scarce. Although electroencephalographic (EEG) recordings from the avian hippocampus homologue revealed clear theta rhythm in awake homing pigeons (16), SWRs were not reported in this study. Other reports investigated sleep EEG activity in birds (17–19), but SWRs were not directly investigated (20).

We reexamined the existence of avian SWRs by recording the LFP from two avian species which diverged 80 million years ago (Fig. 1A). Whereas the chicken represents a basal avian species of neoagnaths, the zebra finch represents an evolutionarily younger avian species. Motivated by the findings of (13) we searched for avian SWRs not in the avian hippocampus homologue (the avian hippocampal formation) but in the avian DVR (Fig. 1B).

**Figure 1.**
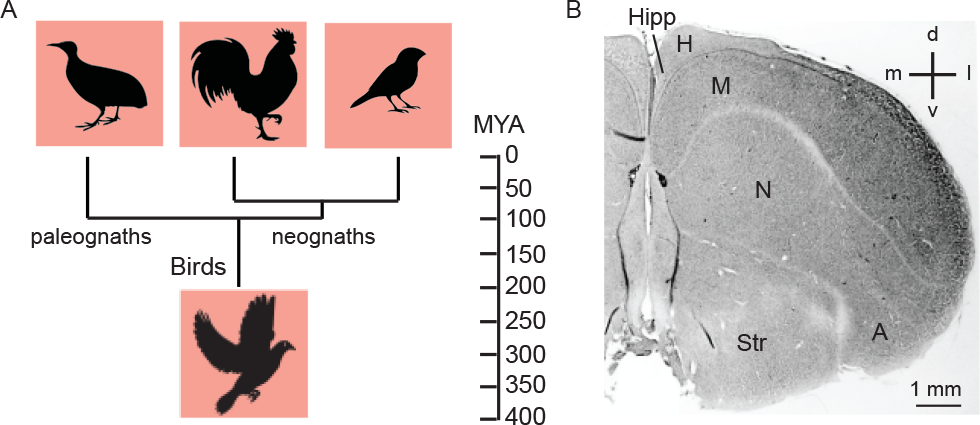
Evolution of modern birds. **(A)** Cladogram of avian evolution. The two living clades of birds are the paleoagnaths and the neoagnaths. The paleognaths, birds such as the tinamou, ostrich, and kiwi, split from the neognaths around 100 million years ago (MYA). Among the neoagnaths, the chickens split from finches around 80 MYA (7, 8). **(B)** Coronal section through the zebra finch brain. Main pallial features are indicated. The avian DVR is composed of the mesopallium (M), the nidopallium (N), and the arcopallium (A)(9, 10). The avian hippocampus homologue (Hipp) is located medially. H, Hyperpallium; Str, Striatum; dorsal; v, ventral; m, medial; l, lateral.

## Results

### Avian SWRs occur during sleep

We used 16-channel silicon probes implanted chronically into the zebra finch DVR (mesopallium; Fig. S1) to measure the brain activity during sleep. After recovery from surgery, we recorded extracellular electrical activity [LFP, multi- and single-unit activity] continuously over 14 to 16 hours centered on the middle of the night. We combined electrophysiological recording with behavioral monitoring to observe the sleeping state of the animals, and recordings from individual animals were repeated for 5 to 7 consecutive days.

LFP recordings revealed rich brain dynamics during sleep (Fig. 2A). A prominent feature of the LFP was the occurrence of negative deflections of varying amplitude (Fig. 2B) separated by short segments of low amplitude, broadband activity (Fig. 2C). The negative deflections were often (18.01±1.78%) associated with a high frequency oscillation (Fig. 2D-E). These avian sharp wave-ripples (SWRs) were highly similar to reptilian SWRs recorded in the reptilian DVR (13), and by analogy we refer to the negative deflections as “sharp waves” and the high frequency oscillation as “ripples.”

**Figure 2.**
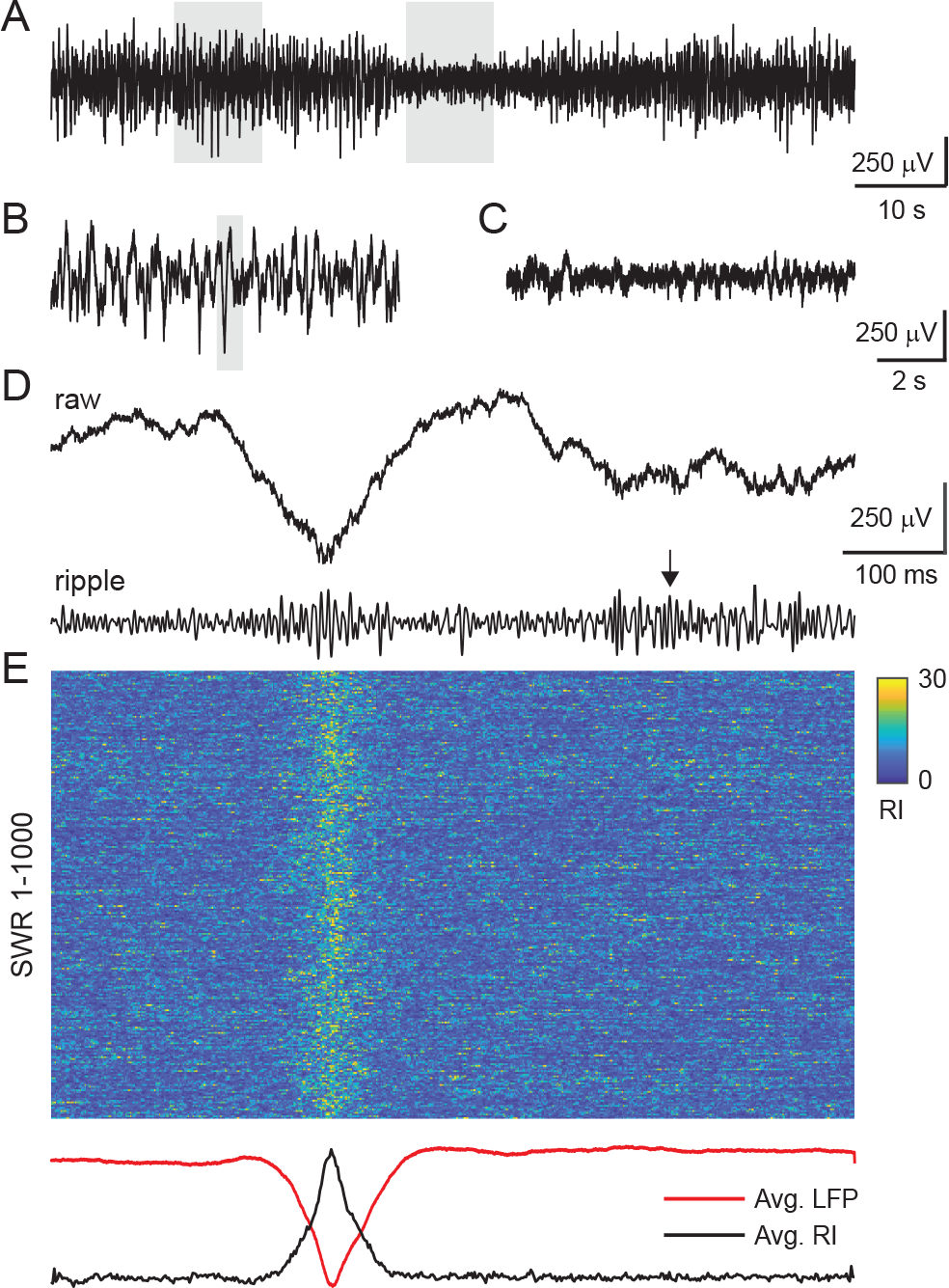
Features of LFP recorded during sleep in the zebra finch. **(A)** 90 s of raw LFP (black) recorded during sleep. Large amplitude negative deflections (gray box, left; expanded in B) are separated by low amplitude, broadband activity (gray box, right; expanded in C). **(B)** Stereotypical negative LFP deflections. **(C)** Broadband LFP that lacks large amplitude negative deflections. **(D)** Close up view of the shaded deflection in (C). The same single trace is shown below after filtering the data in the ripple band (80-300 Hz). Note how ripples are present at the trough of the negative deflection, but also occur afterwards (black arrow). **(E)** One thousand successive SWRs aligned, for each trace, on the peak negative LFP waveform. Averages for all 1000 matching ripple intensity (RI, black) and LFP (red) traces are shown below.

### Switching between high and low delta phases

During sleep, phases of high δ power (1-4 Hz) coincided with periods of sustained SWR activity. We computed the ratio of δ to γ power (25-140 Hz; Fig. 3A) to track SWR activity during sleep, as these spectral bands were shown to be prominent during zebra finch sleep (Fig. 3B; (21)).

**Figure 3.**
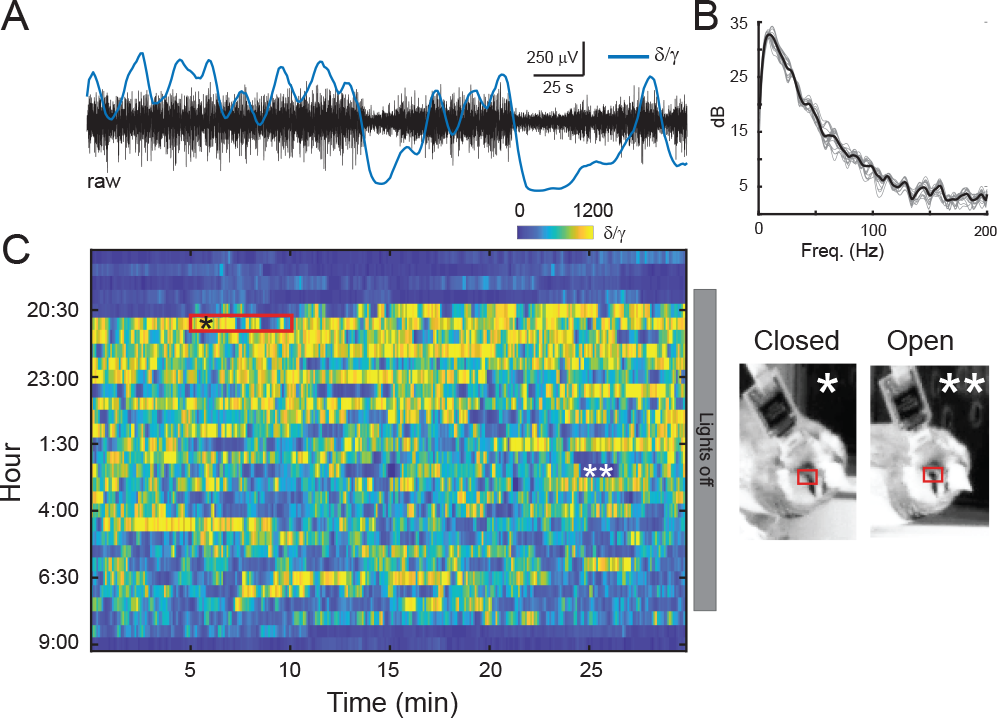
SWR activity during sleep coincides with high delta power. **(A)** 5 minutes of raw LFP (black) recorded during sleep. Data corresponds to red box in (C). The LFP is composed of distinct states, as evident in the *δ*/*γ* ratio (blue line), where large amplitude LFP coincides with high *δ* power. **(B)** Power spectrum calculated on the data in (A) reveals peaks in low frequency bands (1-25 Hz). Gray lines indicated the power spectrum calculated for 20 individual 10s LFP segments; the black line is the mean. **(C)** Epochs of high and low *δ* power begin shortly after lights turn off and continue throughout the night until just after the lights turn on. Periods of high *δ* power coincide with eye closure (*, red box), whereas periods of low *δ* power sometimes correspond to eye opening (**, red box). (The brightness of the images was increased to make the eyes more visible.)

Beginning shortly after lights off, the brain activity switched through epochs of high and low δ power through-out the night until just after the lights turn on (Fig. 3C; Fig. S2). No clear periodicity in state switching was apparent over nights in birds, which is in line with electrophysiological evidence from several avian species reporting variable durations of REM and SWS phases (17, 21–23).

Sharp waves occurred both with and without ripples, and ripples occasionally occurred without sharp waves (Fig. S3A). We detected sharp waves and ripples separately in a two-step process by 1) detecting the sharp waves on the inverted, low pass filtered (< 40 Hz) LFP (Fig. S3B) and 2) by detecting the ripples (Fig. S3C) on the rectified and squared ripple-band filtered (80-300 Hz) LFP (Fig. S3D). Peaks were detected as threshold crossings, and thresholds were defined separately for the sharp waves and ripples (see Methods section for more details). SWRs were defined as sharp waves that coincided with a ripple that occurred within a 60 ms window centered on the trough of the negative deflection, and sharp waves (SW) were defined as sharp waves that occurred without ripples (Fig. S3E-H). SWRs comprised 16.63 - 23.81% of all detected SWs. Unless explicitly stated otherwise, all of our analysis focused on SWRs.

### Avian SWR amplitudes and durations during sleep

We examined SWR amplitude, duration, and rate over the course of a night of natural sleep (Fig. 4). Behavioral analysis revealed that periods of movement (Fig. 4A, gray peaks) were interrupted by short bouts of quiescence before the lights turned off). During the 12 hours of light off (20:00-8:00 hr), a total of 14,820 SWRs were detected. The median SWR amplitude was 147.91±0.46 μV, and the median SWR duration was 53.98±0.15 ms.

**Figure 4.**
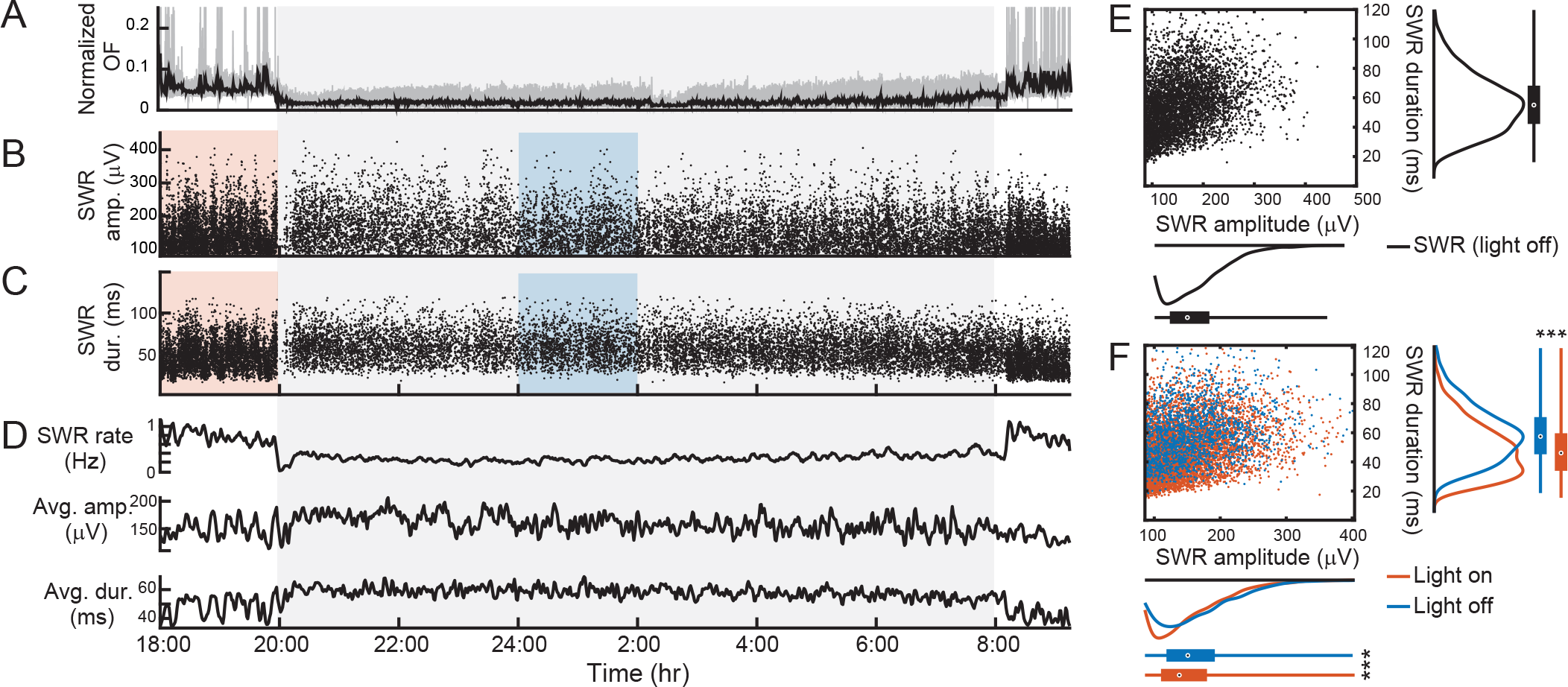
SWR amplitude, duration, and rate during a night of sleep. **(A)** Normalized optic flow estimation reveals bouts of movement during lights on (gray peaks) that subside during lights off (gray shading; 20:00-8:00). Black line, smoothed optic flow (60 s). **(B)** SWR amplitude as a function of time. Each dot represents a single SWR. **(C)** SWR duration as a function of time. Same figure convention as in A. **(D)** Average SWR rate (top), average SWR amplitude (middle), and average SWR duration (bottom). Note how the SWR rate decreases to less than 0.3 Hz during the period of lights off. Similarly, the average SWR duration increases during the period of lights off. **(E)** Scatter plot indicates the SWR amplitude versus duration for 5000 randomly selected SWRs detected during lights off. The marginal distributions are plotted as histograms for the SWR durations (right) and amplitudes (bottom). Bar plot bottom and top edges represent the 25th and 75th percentile, respectively, and the middle dot represents the median. Whisker extends to all points outside of the 25-75 percentile range. **(F)** Scatter plot of SWRs detected during 2 hours of lights on (orange rectangle in A, B; orange dots) and 2 hours of lights off (blue rectangle in A, B; blue dots). Figure conventions same as in E. SWR distributions during lights on (orange) and lights off (blue) were significantly different (***) with respect to amplitude and width (p < 0.001; Wilcoxon rank-sum test).

We compared SWRs detected during two hours of lights on (18:00-20:00 hr; Fig. 4B-C, orange shading) to SWRs detected during two hours of lights off (24:00-2:00 hr; Fig. 4B-C, blue shading). Although the SWR rate was reduced during sleep compared to awake periods (Fig. 4D, top trace), SWR amplitude and duration were significantly different (p < 0.001; Wilcoxon rank-sum test). SWR amplitudes during sleep were significantly larger (median SWR amplitude (light off): 150.81±1.32 μV compared to median SWR amplitude (light on): 137.94±0.71 μV) and SWR durations during sleep were significantly longer (median SWR duration (light off): 57.48±0.44 ms compared to median SWR duration (light on): 46.12±0.24 ms.

To compare SWR statistics across nights, we focused on the SWRs that occurred during lights out (20:00-8:00 hr) over five consecutive sleep recordings (N2-N6) and examined SWR rates, amplitudes, and durations. SWR detections for each night varied (median SWR n = 14,820± 2,254; n = 8,937 SWRs (N6) to 21,162 SWRs (N2)). We calculated SWR rates across nights by binning all detected SWRs into 60 s bins and calculating the SWR rate for each bin (Fig. 5A). Median SWR rates were low and ranged from 0.15±0.01 Hz (N2 and N6) to 0.42±0.01 Hz (N2). We compared SWR am plitudes and durations across consecutive nights for all detected SWRs. Median SWR amplitudes ranged from 146.97 ±0.43 μV (N4) to 165.16 ±0.67 μV (N6); Fig. 5B). Median SWR durations ranged from 48.87 ±0.20 ms (N6) to 56.22 ±0.12 ms (N2; Fig. 5C). We found a small but significant correlation between SWR amplitude and duration (R = 0.25, p <0.001; Pearson’s correlation coefficient).

**Figure 5.**
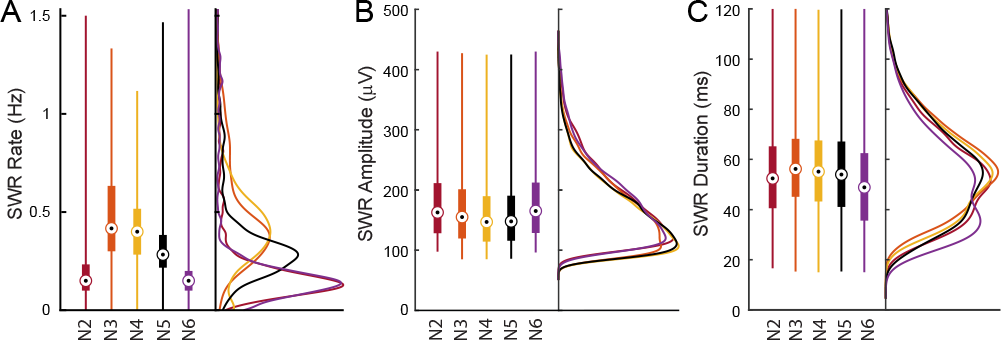
SWR amplitude, duration, and rate for consecutive nights of sleep. **(A)** Comparison of SWR rates calculated on 60-s bins for consecutive recording nights (20:00-8:00 hr). Consecutive nights (N2-N6) represented as different colors; data from N5 is presented in Fig. 4). Box bottom and top edges represent the 25th and 75th percentile, respectively, and the middle dot represents the median. Whisker extends to all points outside of the 25-75 percentile range. The corresponding marginal distributions are plotted as histograms to the right of the box plots. **(B)** Comparison of SWR amplitudes for consecutive recording nights. Figure conventions same as in panel A. **(C)** Comparison of SWR durations for consecutive recording nights. Figure conventions same as in panel B.

### SWRs in anesthetized birds resemble SWRs during natural sleep

In order to investigate the avian SWRs in greater details, we used 16-channel silicon probes and tungsten electrodes to record throughout the depth of the avian DVR in head-fixed, anesthetized zebra finches (n = 2) and chickens (n = 3). A metal electrode was used to record SWRs in chicken 1 (C1; Fig. 6, red trace) and chicken 2 (C2; Fig. 6, orange trace). We used an acute, 16-channel silicon probe to record SWRs in chicken 3 (Fig. 6, yellow trace), zebra finch 1 (ZF1; Fig. 6, purple trace) and zebra finch 2 (ZF2; Fig. 6, green trace). We used a chronically-implanted, 16-channel silicon probe to record from zebra finch 3 under anesthesia (ZF3, Fig. 6, blue trace) and during sleep (Fig. 6, black trace). We typically recorded 20-30 minutes at depths ranging from 1.0-4.0 mm in each anesthetized animal.

**Figure 6.**
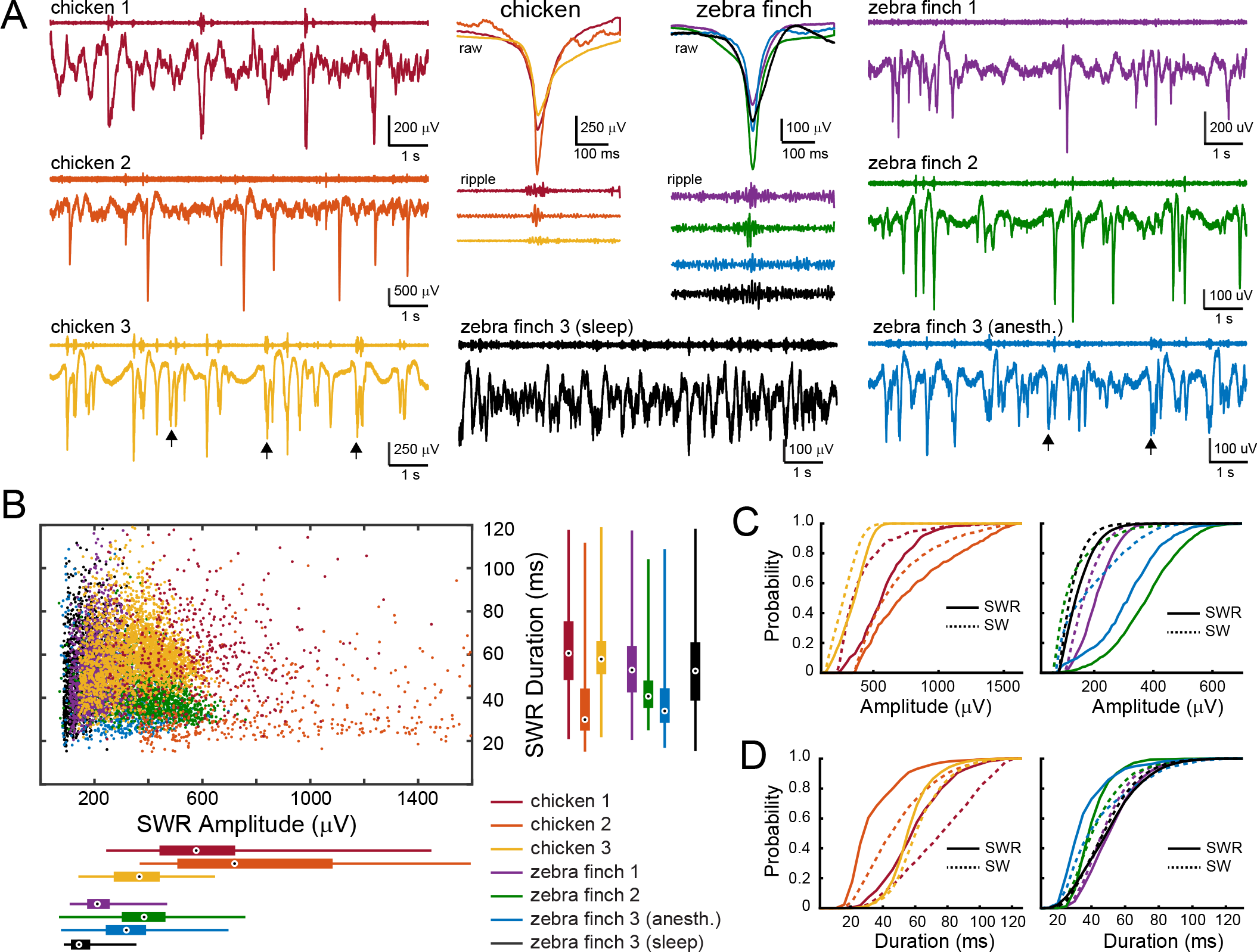
SWR across avian species are similar under anesthesia. **(A)** SWR comparison of 10 s of LFP recorded from 3 anesthetized chickens (left panels, warm colors) and data recorded from 3 anesthetized zebra finches (right panels, cool colors). For each trace, the ripple band (upper plot) and the raw LFP (lower plot) are shown. Complexes of multiple SWRs are indicated as black arrows. The lower middle black trace shows the natural sleep (non-anesthetized) data for zebra finch 3; the anesthetized data for zebra finch 3 was recorded during the electrode implantation surgery. The middle panels depict the average raw traces of 250 SWRs for chickens (left) and zebra finches (right); the lower traces depict the mean ripple-filtered LFP (ripple band: 80-300 Hz) for the corresponding color-coded SWRs. **(B)** Scatter-plot depicts SWR amplitude and widths for SWRs detected from recordings depicted in A. Color-coded bar-plots summarize the SWR durations (right panel) and the SWR amplitudes (lower panel). Box bottom and top edges represent the 25th and 75th percentile, respectively, and the middle dot represents the median. Whisker extends to all points outside of the 25-75 percentile range. For the sake of clarity, only 1000 randomly selected SWRs are plotted for zebra finch 3 (sleep). **(C)** Cumulative distribution plots indicates the amplitude differences for SWRs (solid lines) versus SW (dashed lines) for the chickens (left plot) and the zebra finches (right plot). In all cases, SWR amplitudes were significantly different from the SW amplitude within each animal (p < 0.001, Wilcoxon rank-sum test). **(D)** Cumulative distribution plots indicate the duration differences for SWRs versus SW. Convention same as in C. In all cases, SWR amplitudes were significantly different from the SW amplitude within each animal (p < 0.001; Wilcoxon rank-sum test).

Similar to SWRs recorded during natural sleep, SWRs recorded under anesthesia were characterized by large amplitude negative deflections combined with a high frequency oscillation (Fig. 6A). SWRs occasionally appeared as complexes of multiple sharp waves under anesthesia in both chickens and zebra finches (Fig. 6, black arrow). SWRs in anesthetized zebra finches were larger in amplitude compared to natural sleep (ZF3 median SWR amplitude under anesthesia was 318.42±5.06 μV, n = 538 SWRs compared to ±0.46 μV, n = 14,820 SWRs during sleep). Similarly, SWRs in anesthetized zebra finches were shorter in duration compared to natural sleep (ZF3 median SWR duration under anesthesia was 33.90±0.66 ms compared to 53.98±0.15 ms during sleep). SWRs recorded in chickens using metal electrodes (C1 and C2) tended to be larger than those recorded using silicon probes (Fig. 6B; red and orange box-plots compared to others).

Distributions of SWRs compared to SWs without ripples were significantly different with respect to amplitude (Fig. 6C) and duration (Fig. 6D) for both chickens and zebra finches (p<0.001; Wilcoxon rank-sum test).

### LFP recorded across electrode sites are highly correlated under anesthesia

Under anesthesia, the LFP recorded across the 16 electrode sites were highly similar (Fig. 7A, C, E). We quantified the correlations between electrode sites by calculating Pearson’s correlation coefficient on 10 s traces of LFP for each pairwise channel comparison. Correlations were very high for neighboring sites, and clusters of highly-correlated sites were apparent (Fig. 7B, D, F; black dotted outline). Highly correlated clusters corresponded to anatomical ranges of approximately 0.7 mm in chicken (Fig. 7B, upper and lower clusters) to 0.3-0.9 mm in zebra finch (Fig. 7D, upper and central clusters). We compared correlations under anesthesia and sleep in the same animal (Fig. 7E-H). Correlations between LFP traces were weaker across electrode sites during natural sleep (Fig. 7H) compared to the anesthetized state (Fig. 7F).

**Figure 7.**
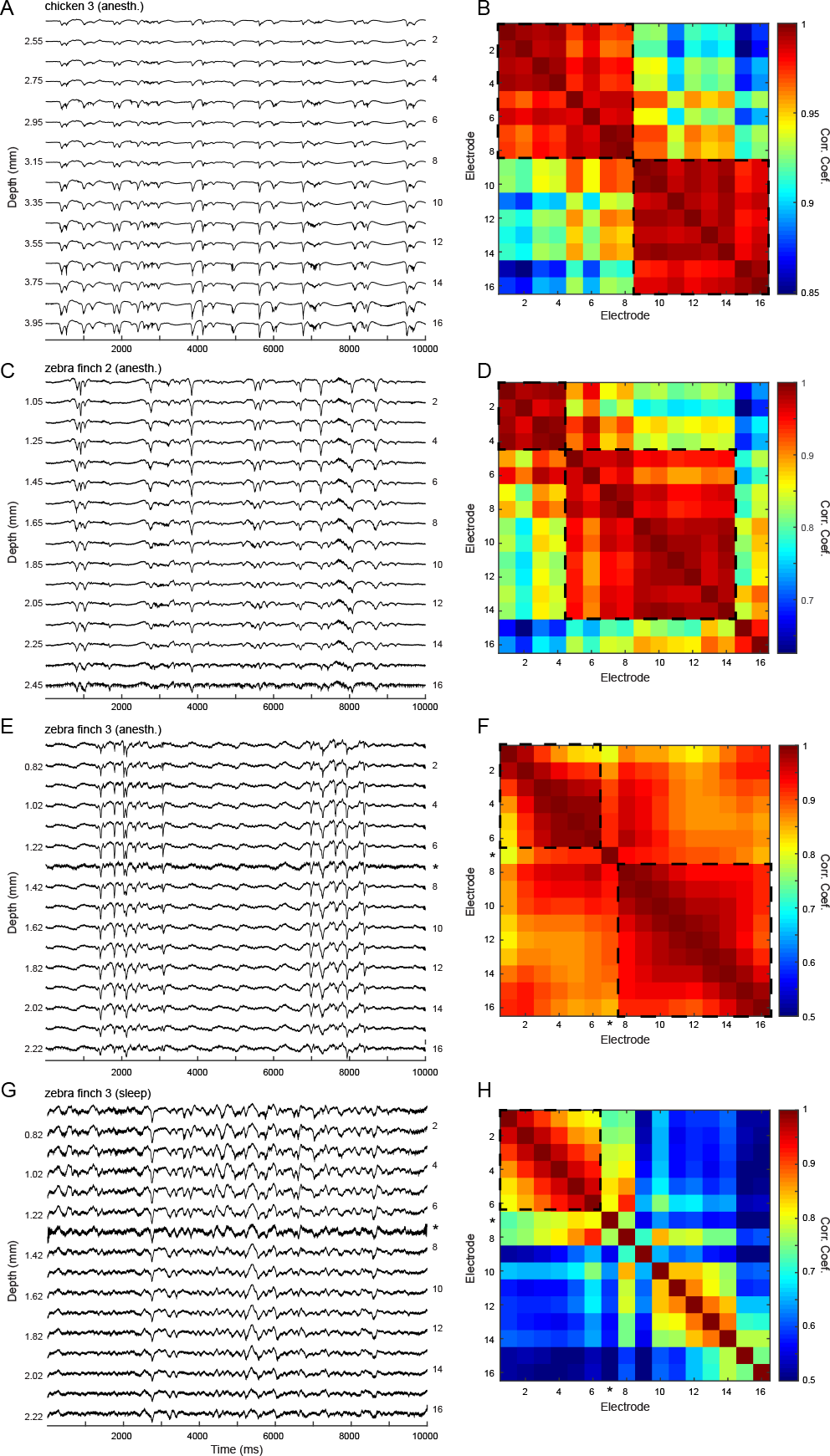
Strong LFP synchrony exist across electrode sites under anesthesia. **(A)** 10 s of raw LFP recorded in an anesthetized chicken (C3). Electrode depth is indicated on the left, electrode number is indicated on the right. Note how the largest amplitude signals are recorded on the deeper electrodes > 3.25 mm. Vertical scale: 1200 *μ*V. **(B)** Matrix indicates correlations between each pairwise comparison of the traces in A (Pearson’s correlation coefficient). Warm colors indicated large R values, cool colors indicated small R values. Note strong correlations for electrodes 1-8 and 9-16 (dotted black outline). **(C)** 10 s of raw LFP recorded in an anesthetized zebra finch (ZF2). Note how the largest amplitude signals are recorded on the superficial electrodes. Figure conventions same as in A. Vertical scale: 600 *μ*V. **(D)** Matrix indicates correlations between each pairwise comparison of electrode channels in C. Figure conventions same as in B. **(E)** 10 s of raw LFP recorded in chronically implanted zebra finch under anesthesia (ZF3). Figure conventions same as in A. Asterisk indicates a defective site. Note how the largest amplitude signals are recorded on the superficial electrodes. Vertical scale: 500 *μ*V. **(F)** Matrix indicates correlations between each pairwise comparison of electrode channels in E. Asterisk indicates a defective site. **(G)** 10 s of raw LFP recorded in a zebra finch during natural sleep (ZF3: N3). Figure conventions same as in A. **(H)** Matrix indicates correlations between each pairwise comparison of electrode channels in G. Figure conventions same as in B. Note how electrode sites are less correlated during sleep than under anesthesia (comparison of F and H, same color scale).

### Stereotyped spike sequences during SWR events

An increase in spiking activity coincided with the occurrence of SWRs: across consecutive SWRs, the population spiking activity was tightly locked to the SWR, and spiking activity outside of SWR events was low (Fig. 8). However, spiking also occurred during ripple events not associated with a SW (Fig. 8A, black arrow). For both chickens and zebra finches, stereotyped spiking responses corresponded to the electrode sites with the highest amplitude SWRs (Fig. 8A, lower electrode sites; Fig. 8B, upper electrode sites). Importantly, not every neuron participated in every SWR, indicating that different neural ensembles may participate in different SWR events.

**Figure 8.**
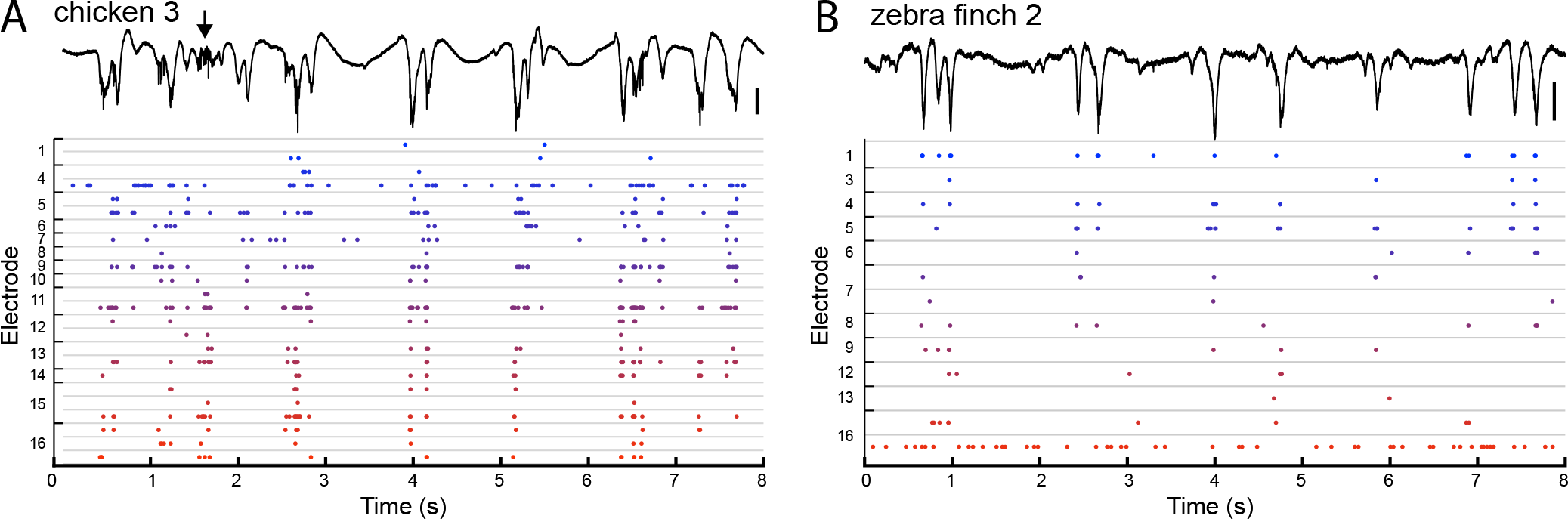
Spiking sequences across consecutive SWRs. **(A)** Top panel: C3 LFP containing multiple SWR events. Same recording as Fig. 7A. Black arrow indicates a ripple event that occurred without a corresponding SW. Vertical scale bar, 250 μV. Lower panel: Sorted spikes that correspond to the trace in the upper plot. Each dot indicates a spike. Thin gray lines indicated distinct spike clusters. In some cases, several spike clusters were isolated from the same electrode site (indicated on the left). Spiking across the population was aligned to the SWR events, and stereotyped spiking responses were located on the deeper electrode sites with the largest amplitude SWRs. Also note how increased firing was locked to the the ripple event. **(B)** Top panel: ZF 2 LFP containing multiple SWR events. Same recording as Fig. 7C. Vertical scale bar, 200 μV. Lower panel: sorted spikes that correspond to the trace in the upper plot. Firing rates were lower compared to chicken, but stereotyped responses were also located on the electrode sites with the largest amplitude SWRs.

**Figure 9.**
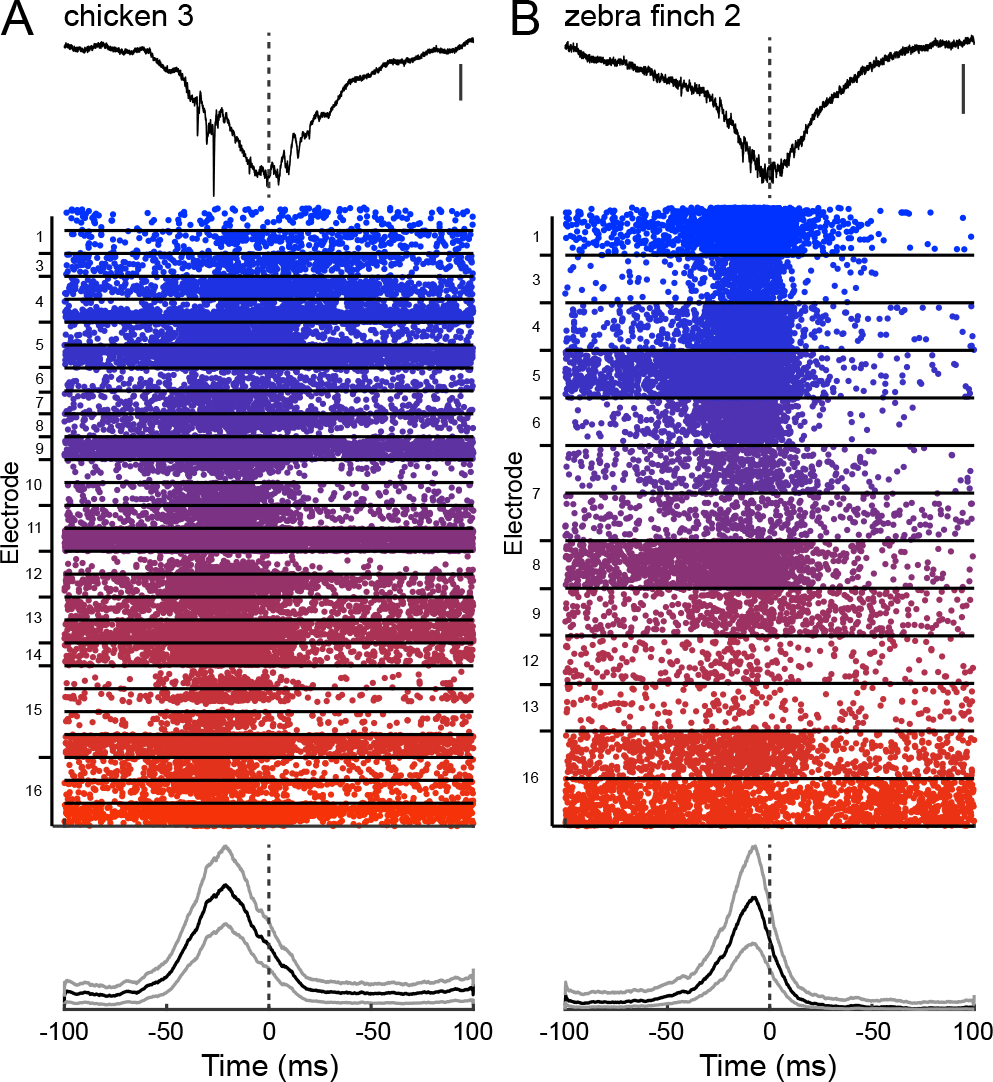
Mean population spiking activity precedes negative SWR deflection. **(A)** Upper panel: LFP of exemplary SWR in C3. 200 ms window is centered on the negative SWR deflection (dotted line). Same recording as Fig. 8A. Vertical scale bar, 200 μV. Middle panel: Spike rasters aligned to all SWR detections (n = 864). Each dot indicates a spike. 27 distinct clusters are demarcated with black horizontal lines. Some clusters were isolated from the same electrode site (indicated to the left). Lower panel: Mean firing response averaged over all clusters (5 ms smooth window). Black line indicates the mean, gray lines indicates the mean ± s.e.m. Dotted black line indicates the trough of the negative deflection. Note how mean population firing precedes the negative deflection in time (peak = −20.4 ms). **(B)** Upper panel: LFP of exemplary SWR in ZF2. Same recording as Fig. 8B. Vertical scale bar, 200 μV. Middle panel: Spike rasters aligned to all SWR detections (n = 1,426). 13 distinct clusters are demarcated with black horizontal lines. Figure conventions same as in A. Note how mean population firing also precedes the negative deflection (peak = −7.3 ms).

### Population spiking precedes negative SW deflection

We aligned spiking activity to the trough of the SWR deflection (Fig. 8A, B). In both chickens and zebra finches, population spiking activity increased prior to the trough of the negative SWR deflection (chicken peak = −20.4 ms; zebra finch peak = −7.3 ms). This peak in firing preceding the negative SWR deflection is not accounted for by the variation of SWRs durations (C2 mean SWR duration = 62.09±0.52 ms; ZF2 mean SWR duration = 42.899±0.28 ms).

## Discussion

In this paper, we have reported the first evidence for avian SWRs recorded from the DVR during sleep and under anesthesia. SWRs were highly similar in chickens and zebra finches. Similar events have also been described in anesthetized pigeons (24). This evidence for avian SWRs follows similar evidence in sleeping reptiles (13, 14) and suggests that the evolutionary origin of large-amplitude, highly synchronous SWR events may have already been present in the stem-amniote ancestor to birds, reptiles, and mammals.

Mammals, reptiles, and birds are descendants of a now extinct, stem-amniote ancestor. These amniotes were the ancestors of synapsids (mammals) and reptiles and birds (sauropsids). The sauropsids are a diverse group of egg-laying amniotes that includes Squamata (lizards and snakes) and a group comprising both the turtles and the Archosauria (dinosaurs, modern birds, and crocodilians) (25).

The brain organization of amniotes varies widely between mammals, reptiles, and birds. In mammals, the pallium has a laminar organization that spans the medial pallium (the hippocampus), the dosal pallium (the isocortex) and the lateral pallium (the olfactory cortex) (26).

In reptiles, the dorsalmost and medial aspects of the pallium make up a primitive 3-layered structure that differs from that of the more basal amphibians. The lateral and ventral pallium, on the other hand, is a relatively large structure that bulges inside the ventricular cavity, which is termed the DVR (26). The anterior DVR consists of spatially segregated neuronal types specialized in processing visual, auditory, or somatosensory stimuli (27), whereas the posterior DVR is homologous with the mammalian claustrum and amygdala (28).

In birds, the DVR has become severely hypertrophied, and can be subdivided into (i) a nidopallium (originating from the ventral pallium and receiving visual and auditory mesencephalic sensory input) and (ii) a mesopallium (originating from the lateral pallium) (29). These structures are morphologically different from that of the mammals, in that there is no laminar organization (26). Although the avian DVR is developmentally different from the mammalian hippocampus, its placement as a nexus of sensory input within the avian telencephalon might make DVR networks well-suited for a role in learning and memory. Further work is thus needed to clarify the relationship between the DVR, a dominant part of the avian forebrain, and its potential mammalian equivalent.

The SWRs in birds resemble those identified in rodent hippocampal CA1 (30) both in terms of the large amplitude, negative deflection and the corresponding high-frequency ripple. In contrast to mammalian sharp wave and ripple events, which are spatially segregated in the stratum radiatum and the CA1 pyramidal layer (3), avian SWs and ripples - like reptilian SWRs (13) - are superimposed sharp wave-ripple events that are recorded from the same spatial location.

Similar to mammalian SWRs, avian SWRs were highly synchronous events in birds that spanned up to > 1 mm of tissue along the electrode tract. The largest amplitude SWR events were usually localized to a specific area of the electrode tract, suggesting that there may be a single anatomical source of SWRs in avian DVR; however, more anatomical work is necessary to determine the exact location and circuitry underlying SWR generation. Correlation analysis found that clusters of highly correlated responses were apparent, and furthermore, that these responses were more strongly correlated under anesthesia compared to natural sleep.

Analysis of the spiking activity associated with SWR events under anesthesia found that spiking activity across the population was tightly locked to SWR and ripple events, and spiking activity outside of the SWR events was low. On average, the population spiking activity preceded the negative SWR deflection by 7-20 ms. This evidence suggests that there could be a gradual buildup of activity that triggers the SW event.

In the mammalian hippocampus, SWRs initiate when firing in a set of spontaneously active pyramidal cells triggers a gradual, exponential buildup of activity in the recurrently connected CA3 network (31). This tonic excitation drives reciprocally connected parvalbumin-positive interneurons, which gives rise to phase-locked, ripple-frequency spiking. More work needs to be done to investigate the subthreshold activity underlying the avian SWR activity, and specifically to investigate the IPSCs and EPSCs that correspond to the SWR events, and the role of excitatory and inhibitory neurons in these events in general.

Finally, preliminary evidence suggests that neurons may fire in specific sequences during consecutive SWR events. Specifically, not every neuron participated in every SWR event, which suggests that different neuronal ensembles may participate in different SWR events. Although more work needs to be done to quantify this effect, this evidence is remarkably similar to CA1 firing patterns during SWS ripple events in rats (32).

Although avian SWRs resemble mammalian hippocampal SWRs in terms of shape, frequency content, network synchrony, and population spiking patterns, the major detail that is still lacking is the behavioral learning component. Do avians SWRs have anything to do with learning? One of the most fascinating features of hippocampal place cells is their reactivation associated with SWR events during SWS: cells that fire together when the animal occupied particular locations in the environment exhibit an increased tendency to fire together during subsequent sleep (11). This role in memory consolidation still needs to be addressed for avian SWRs events, and songbird vocal learning promises to be a useful tool for the study of memory consolidation events during sleep (33–35).

## Methods

### Experimental animals

Experiments were conducted on 3 adult zebra finches (*Taeniopygia guttata*) and 3 juvenile chickens (*Gallus gallus domesticus)*.

#### Zebra finches

Zebra finches (2 females, 1 male; ages 597-1055 days post hatch (dph)) were obtained from the Free University Berlin in 2017 and housed on site in volieres. Birds were kept on a 12:12-hr light:dark cycle. Finches received seed mix, millet, sepia bone, and water *ad libitum* and were given fresh salad and cooked egg supplements once a week.

#### Chickens

Fertilized chicken eggs were obtained from Prof. Dr. Benjamin Schusser, Reproductive Biology, TUM School of Life Sciences-Weihenstephan and reared on site. After hatching, young chickens (2 males, 1 female; ages 20-71 dph) were kept in social groups in small arenas on a 12:12 hr light:dark cycle and had access to sand baths, perches, water and food *ad libitum*.

All experiments were carried out in accordance with Oberbayern, German, and European laws on animal experimentation (Approval #ROB-55.2-2532.Vet_02-18-108: J.M.O.; #ROB-55.2-2532-Vet_02-18-154: H.L.). The data presented in this paper originate from 6 animals, 5 nights (80 hours) of chronic recording, and 34 hours of acute recording.

### Behavior and video acquisition

Zebra finches were acclimated to the recording chamber for at least 2 weeks prior to the experiments. Experimental animals were kept together with a non-experimental female bird to prevent social isolation. Birds were separated from each other with a clear Plexiglas panel, such that animals could hear and see each other but could not interact physically.

Finches received seed mix, millet, sepia bone, and water *ad libitum*. The recording chamber (total inner measurements: 120 cm × 50 cm × 50 cm) was equipped with LED lights, a UV bird light (Bird Systems, Germany), an infrared (IR) LED panel (850 nm), and had continuous air circulation. Video footage of the experimental animal was acquired with a near-IR sensitive camera (acA1300-60gm, Basler Ag, Germany) with custom-written software (ZR View, Robert Zollner) at a frame-rate of 10 fps, triggered with a pulse generator (Pulse Pal, Sanworks, NY, USA; (36)).

### Anesthesia

Zebra finches were anesthetized with isoflurane (1-3 %) mixed in oxygen generated with a oxygen concentrator (EverFlo OPI, Phillips, Netherlands) and administered using a vaporizer (Isotec 4, Groppler, Germany) through a small tube directly into the beak of the bird. Excess isoflurane was collected and filtered (Scavenger LAS, Groppler, Germany). The body temperature was monitored with a laser thermometer, and maintained at above 39.5°C through the use of an electric heating pad.

Chickens were anesthetized with an initial injection of a mixture of ketamine (40 mg/kg) and xylazine (12 mg/kg) administered intramuscularly (i.m.) into the breast muscle. Afterwards, a mixture of ketamine (20 mg/h/kg) and xylazine (6 mg/h/kg) anesthesia was constantly administered i.m. through a perfusion pump (B. Braun, Germany). The heart rate was constantly monitored by two differential EKG electrodes placed in the breast muscle and contralateral leg muscle (DAM 80, WPI, FL, USA). The body temperature was monitored with a cloacal probe and maintained above 39.5°C through the use of an electric heating pad.

### Electrophysiological recording under anesthesia

Under anesthesia, the experimental animal (2 zebra finches, 3 chickens) was positioned in a small animal stereotaxic frame (Kopf Instruments, CA, USA). The scalp was anesthetized with xylocaine (pump-spray), the feathers were removed with forceps, and the scalp was resected along the midline.

A small craniotomy was drilled above the forebrain with a high speed dental drill (Volvere i7, NSK Europe GmbH, Germany) and a 16-channel silicon probe (NeuroNexus, MI, USA; A1×16: 100 *μ*m pitch, 413 *μ*m2 area for each site; in a single row of 16 contacts), or a 1 MOhm tungsten electrode (Alpha Omega GmbH, Germany) was lowered into the brain. The electrode was connected to a preamplifier (Intan Technologies, RHD2132), which was connected to the data acquisition board with a tether cable (SPI interface cable, Intan Technologies).

Recordings were performed with an Open Ephys acquisition board (OEPS) or a USB Interface Board (RHD2000, Intan Technology). Recordings were grounded and referenced against one of the reference wires. Signals were sampled at 30,000 Hz, wide band filtered (0.1-9987 Hz), and saved in a continuous format using the Open Ephys GUI (37).

### Chronic surgery

Recordings were made from the DVR of a chronically implanted adult male zebra finch. 20-30 minutes before the surgery, the animal was injected with Metamizol (100-150 mg/kg, i.m.) in the breast muscle. After approximately 20 minutes, the animal was anesthetized with isoflurane (1-3%) and prepared as described for “Electrophysiological recording under anesthesia.”

A small craniotomy was drilled above the forebrain and a 16-channel silicon probe was implanted into the DVR (NeuroNexus, A1×16: 100 *μ*m pitch, 177 *μ*m^2^ area for each site; in a single row of 16 contacts). The silicon probe was mounted on a flattened needle secured to a holder. The probe was slowly lowered into the brain stereotaxically to a depth of 2.27 mm. The craniotomy and electrode were covered with silicone adhesive (Kwik-Sil, WPI, FL, USA) and after connecting ground and reference, the skull, craniotomy, and probe were covered with dental cement (Paladur, Henry Schein Dental, Germany).

Following surgery, the animal was released from the stereotaxic frame and administered another dose of Metamizol (100-150 mg/kg, i.m.). The animal remained on a heating pad until full recovery from anesthesia. An antibiotic (Baytril, 1025 mg/kg, i.m.) and an analgesic (Carprofen, 4 mg/kg, i.m.) were administered up to 3 days post-operatively. Following recovery from surgery, the animal returned to entirely normal feeding, sleeping, and singing behaviors as observed before surgery.

### Overnight recordings

Two to three hours before lights off, the animal was connected to the headstage and custom-made, lightweight tether cable and allowed to assume a natural sleep posture on a branch that had been placed on the floor of the chamber. The camera was positioned to allow for the best visualization of the animal during sleep. The animal was allowed to sleep naturally overnight, and was disconnected from the headstage and tether cable shortly after lights turned on. On one occasion, the animal remained connected during the day to obtain awake electrophysiology data for approximately 7 hours. Data from the first night after surgery (N1) was not used in the analysis (Fig. S2A).

### Anatomy

After the period of recording was over, the animal was deeply anesthetized with an overdose of sodium pentobarbitol (250 mg/kg, i.m.) until the corneal reflex disappeared. Afterward, the animal was decapitated, the brain was removed from the skull and post-fixed in 4% paraformaldehyde in PBS, and sectioned on a frozen cryostat (n = 6 animals, 80-100 μm thick, cresyl violet stain).

### Data Analysis

All analysis was performed using Matlab 2019a (MATLAB, MathWorks, USA).

#### Behavioral analysis

The “Lucas-Kanade” method for optic flow estimation (part of the Computer Vision Toolbox for Matlab) was used to determine the optical flow vectors of the video data. A region of interest was defined around the animal’s position in the video. For each frame the horizontal and vertical components of the optical flow were estimated using a 1 frame step. The normalized optic flow was calculated by taking the mean of the absolute value of all the optical flow velocity vectors and dividing by the maximum value.

#### Filtering of LFP

Unless otherwise mentioned, all raw data were band-pass filtered (1-2000 Hz). Occasionally, data were also notch filtered in software to remove 50 Hz electrical noise. δ = 1-4 Hz, γ = 25-140 Hz; ripple band = 80-300 Hz.

#### Spectral analysis of LFP

Voltage traces were band-pass filtered (1-2000 Hz), down sampled (300 Hz) and binned (10s).

The average normalized power spectrum (spectrum in each bin divided by the average over the entire dataset) for each bin was calculated using the Welch method (1 s windows, 50% overlap). The same spectral analysis was used to calculate the δ/γ ratio on 10 s bins (1 s steps) by dividing the mean non-normalized spectrum over frequencies lower than 4 Hz by the mean non-normalized spectrum over frequencies between 25-140 Hz in each bin.

#### SWR detection

SWs and ripples were detected separately in a two-step process. SWs and ripples were generally detected on the electrode site that had the largest amplitude SW and highest ripple content.

For SW detection, the data were bandpass filtered (1-2000 Hz), low pass filtered (<40 Hz) and inverted (such that negative deflections were pointing upwards). For ripple detection, the data were filtered in the ripple band (80-300 Hz), squared, and smoothed (100 ms window). Randomly selected test data (800 s) were used to calculate the thresholds for the SW and ripple detection. Thresholds were set to the value at 90 % of the sorted and filtered SW and ripple test data.

SWs and ripples were detected from consecutive 20 s segments of data (2 s overlap). SWs were detected as peaks that crossed the SW threshold, had a minimum peak width > 10 ms, and an inter-peak distance > 100 ms. Ripples were detected as peaks that crossed the ripple threshold, had a minimum peak width > 10 ms, and an inter-peak distance > 100 ms.

After the initial detection, each SW and ripple was redetected to eliminate double detections and to determine the negative peak of the SW. For SWs, a 200 ms window was centered on the detected SW, and peaks were detected from the inverted band-pass filtered data if they had a minimum peak width > 15 ms and a peak prominence > 5. In case of two peak detections, the smaller of the two peaks was discarded. For ripples, a 200 ms window was centered on the detected ripple, and peaks were detected from the ripple-filtered rectified data if they had a minimum peak width > 15 ms. Very large and small peaks were discarded as outliers in chronic zebra finch recordings: ripples outliers: > 1500 *μ*V 2 and < 115 *μ*V 2; SW outliers: > 430 *μ*V and < 85 *μ*V.

Because the brain activity was generally less dynamic in anesthetized animals, outliers in anesthetized data were usually very large peaks: anesthetized chicken SW outliers: > 1600 *μ*V; anesthetized zebra finch SWs outliers: > 430 *μ*V; anesthetized chickens and zebra finch ripples outliers: (> 1500 *μ*V^2^).

A SWR was defined as a SW that coincided with a ripple that occurred within a 60 ms window centered on the trough of the SW. All SWs that did not meet this criterion were defined as SWs.

#### Ripple intensity (RI)

Ripple intensity was calculated by binning the ripple band (80-300 Hz) filtered data and summing the absolute value of the voltage within each bin (2 ms).

#### Correlation analysis

Correlation analysis was carried out using a pairwise comparisons of the LFP recorded on all electrode channels using Pearson’s correlation coefficient. Specifically, 10 s traces of bandpass filtered LFP (1-2000 Hz) were used in this analysis. All correlations were highly significant (p < 0.001).

#### Spike sorting

Spikes were sorted using JRCLUST (38). Briefly, the continuous Open Ephys files were converted to binary files. Data were filtered using a second order differentiation filter. For each recording site, the spike detection threshold was set to 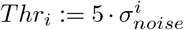, where σ_*noise*_ is the standard deviation of noise distribution for each site *i*. Spike waveform features were extracted using principle component analysis (PCA). Clusters were manually curated into multi-unit or single-unit clusters, and spike times were saved for further analysis.

#### SWR-triggered spiking

All clusters (multi-unit and single-unit) with more than 500 spikes and fewer than 25,000 spikes were analyzed. Spikes from each cluster were aligned within a 200 ms window to the trough of the negative deflection for all detected SWRs (usually detected on the electrode site with the largest signal). The mean firing rate over all clusters was computed and smoothed with a 5 ms window.

#### Statistics

We tested all data for normality using the Liliefors test. Distributions of SWR amplitudes and durations were not normally distributed, therefore we used the the Wilcoxon rank-sum test for equal medians (equivalent to the nonparametric Mann-Whitney U-test). In all cases, uncertainties are represented as the standard error of the mean.

## Acknowledgements

This research was funded by grants from the German Research Foundation (Deutsche Forschungsgemeinschaft; ON 151/1-1) and the Daimler and Benz Foundation to J.M.O. The authors are grateful to B. Seibel, Y.Schwarz, and R. Harpaintner for technical assistance; C. Fink and E. Jochen for help with mechanical design, fabrication, and electronics; L. Hoffman for administrative assistance. Special thanks to Mark Shein-Idelson for his suggestions on the manuscript.

## Author Contributions

J.M.O.conceived of the project, performed experiments, and wrote the manuscript. H.Y and J.M.O. analyzed data. H.L. contributed animals and materials. All authors have discussed analysis, manuscript revisions, and read and approved the final manuscript.

---

**Figure S1.**
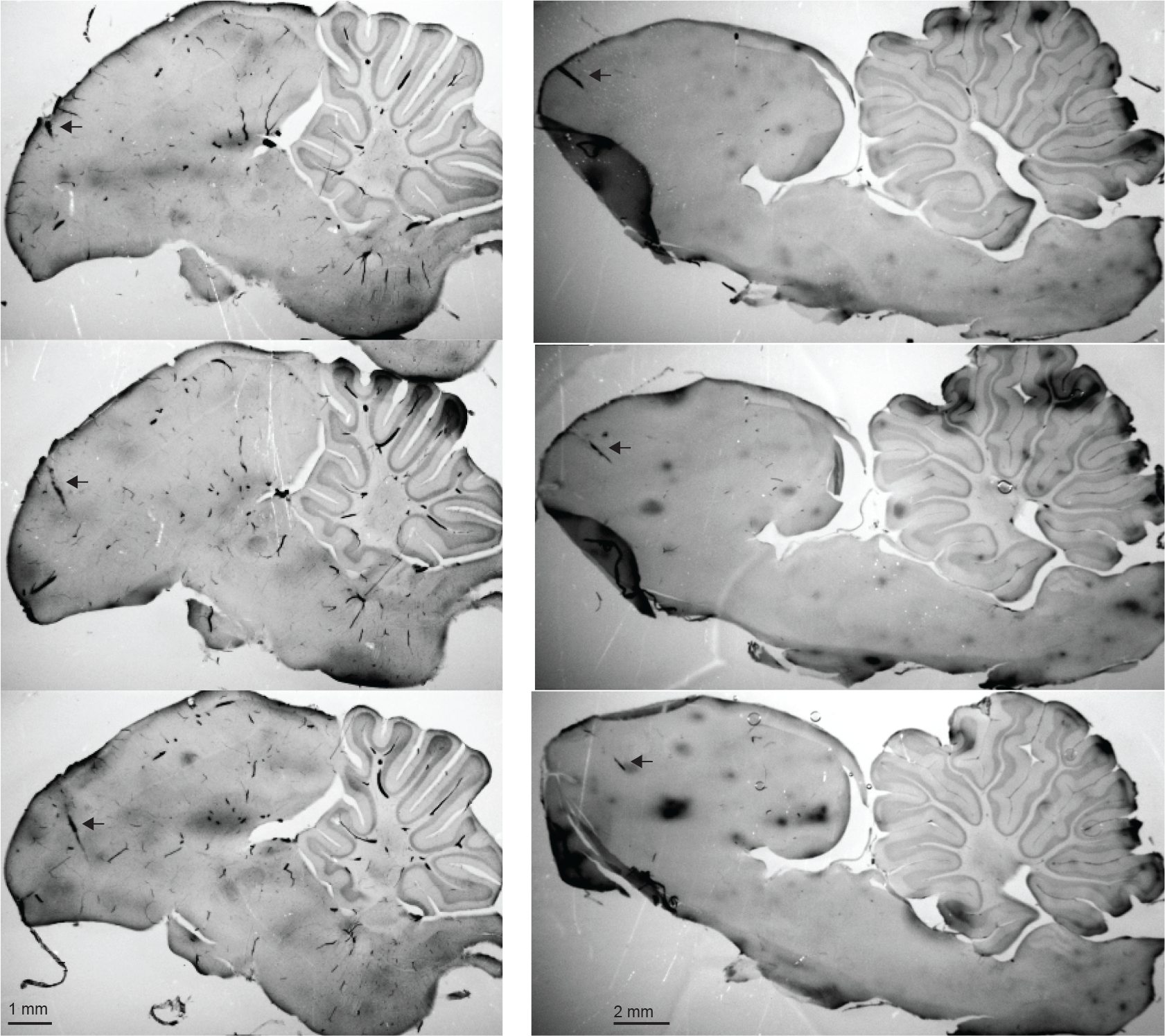
Electrode placement in avian DVR. Three consecutive sagittal sections from the zebra finch (left) and from the chicken (right). Black arrow indicates electrode tract.

**Figure S2.**
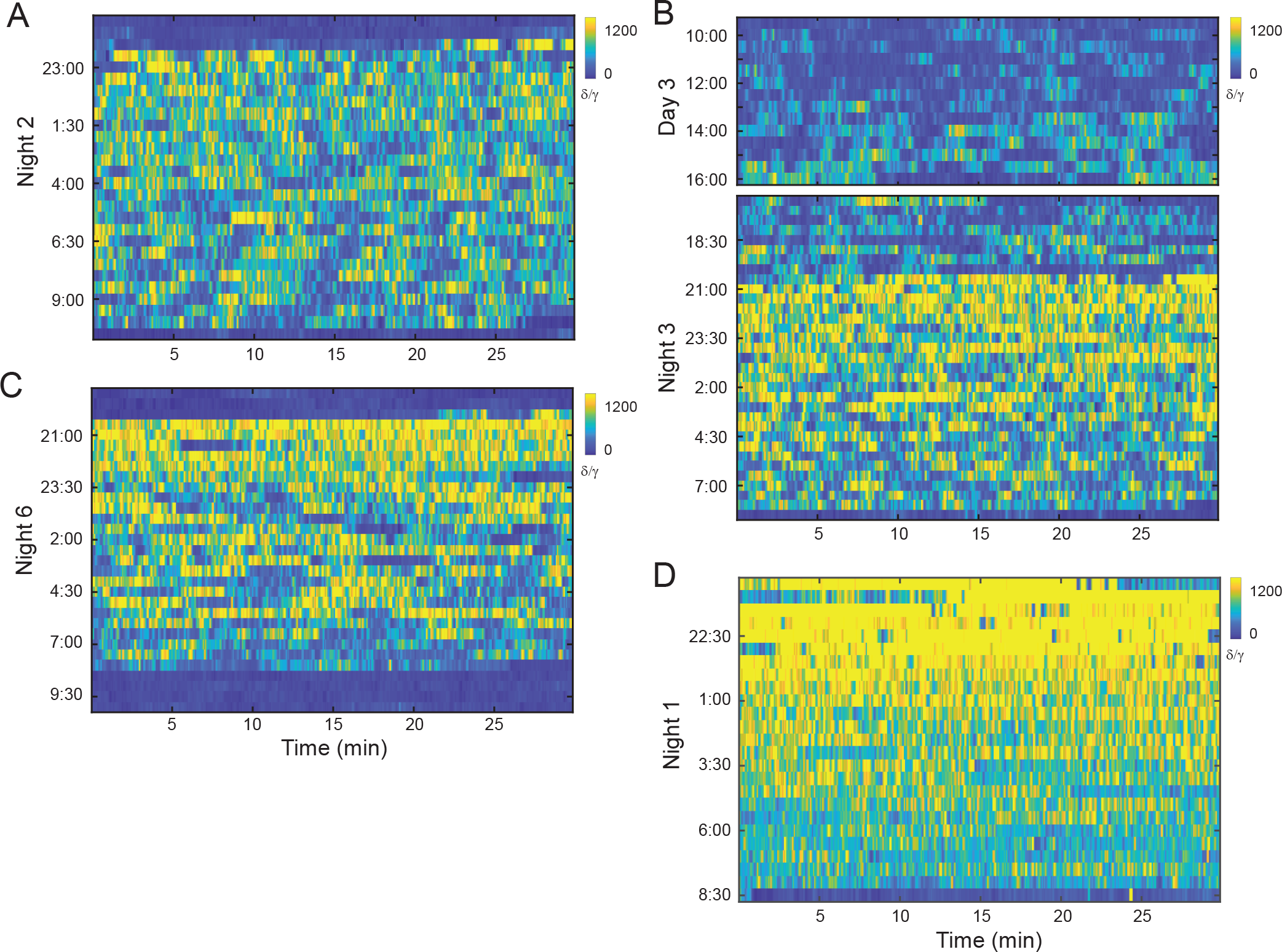
Epochs of high and low δ power are consistent across multiple nights. **(A-C)** Electrophysiological sleep recordings for 3 nights of sleep. As in Fig. 3C, each matrix plots the δ to γ power ratio (measured piecewise over 10s-long data segments, 1 s steps) around night-time (time of night indicated at left). Rows along x represent 30-min segments running continuously (L to R) from top to bottom. Note that for all nights this high/low [δ/γ] alternation starts shortly after light off (20:00 hr), and continues uninterrupted until just after the lights turn on (8:00 hr). Also note the reduced δ to γ power during the 7 hour recording of awake activity during the day (B, upper plot). **(D)** Data from the first night of recording, directly after the surgery, was not used in the analysis, due to the very high δ power present throughout the night, possibly the result of the anesthesia used during the surgery.

**Figure S3.**
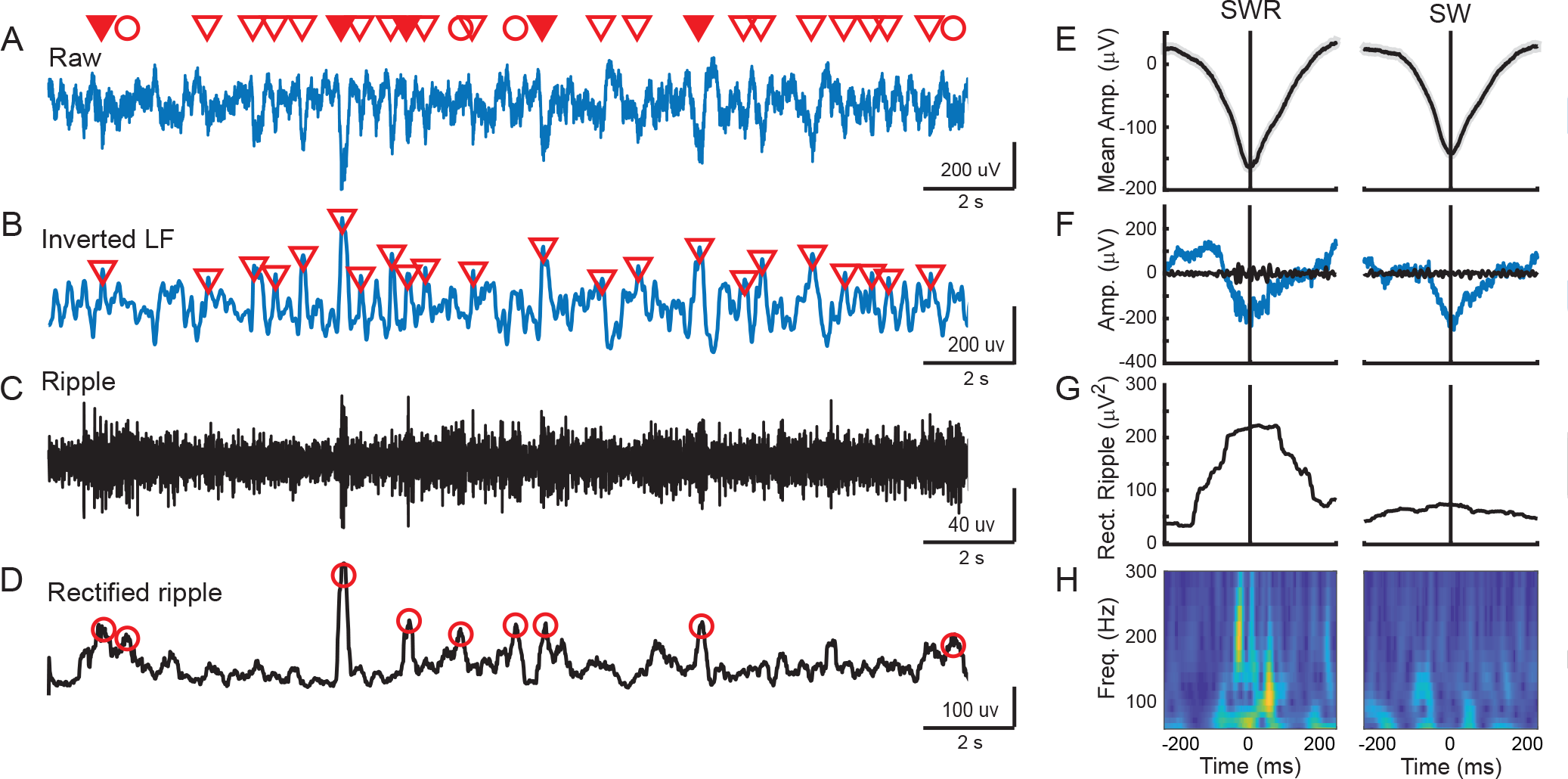
Detection of SWRs, SWs, and Ripples. **(A)** 20 s of raw LFP. Filled triangles indicated SWRs, empty triangles indicated SWs, and circles indicate ripples. **(B)** The same trace in A, low pass filtered (< 40 Hz) and inverted. SWs are indicated as empty triangles. **(C)** The same trace in A, filtered for the ripple band (80-300 Hz). **(D)** The same trace in C, but squared and smoothed (100 ms window). Circles indicate ripples. **(E)** Average of 500 SWRs (left) and average of 500 SW (right). Gray shading indicates the s.e.m. **(F)** Example of raw SWR (left, blue trace) and raw SW (right, blue trace). Black line indicates the ripple filter for the same traces. **(G)** Rectified and smoothed ripple corresponding to traces in F. **(H)** Wavelet transform of raw traces in F. Note how the SWR contains power in the ripple band (left) whereas the power is much reduced for the SW (right). Window displays 60-300 Hz.

